# Rapid and extensive karyotype diversification in haploid clinical *Candida auris* isolates

**DOI:** 10.1101/514745

**Authors:** Gustavo Bravo Ruiz, Zoe K. Ross, Silke Schelenz, Neil A.R. Gow, Alexander Lorenz

**Affiliations:** Institute of Medical Sciences (IMS), University of Aberdeen, Aberdeen, UK; MRC Centre for Medical Mycology, University of Aberdeen, Aberdeen, UK; National Heart and Lung Institute, Imperial College London, London, UK; School of Biosciences, University of Exeter, Exeter, UK

**Author notes:** Correspondence should be addressed to: Alexander Lorenz, Institute of Medical Sciences (IMS), University of Aberdeen, Foresterhill, Aberdeen AB25 2ZD, United Kingdom.

**Keywords:** *Candida auris*, chromosome number, chromosome size, genome size, rRNA genes

## Abstract

*Candida auris* is a newly emerging pathogenic microbe, having been identified as a medically relevant fungus as recently as 2009. It is the most drug-resistant yeast species known to date and its emergence and population structure are unusual. Because of its recent emergence we are largely ignorant about fundamental aspects of its general biology, life cycle, and population dynamics. Here we report the karyotype variability of 26 C. *auris* strains representing the four main clades. We demonstrate that all strains are haploid and have a highly plastic karyotype containing five to seven chromosomes, which can undergo marked alterations within a short time-frame when the fungus is put under genotoxic, heat, or osmotic stress. No simple correlation was found between karyotype pattern, drug resistance, and clade affiliation indicating that karyotype heterogeneity is rapidly evolving. As with other *Candida* species, these marked karyotype differences between isolates are likely to have an important impact on pathogenic traits of *C. auris.*

## Introduction

A current, major concern in medical mycology is the emergence of the multidrug-resistant pathogen *Candida auris.* This species was named according to its first identification as an isolate from the ear canal of a Japanese patient in 2009.^1^ Since then, it has rapidly become a major healthcare threat with hospital outbreaks occurring worldwide.^2-4^ Most *C. auris* isolates also show high levels of resistance to antifungal drugs, including azoles, echinocandins, 5-flucytosine, and polyenes (amphotericin B).^5,6^ C. *auris* is also difficult to eradicate from hospital intensive care wards and as a skin colonizer it can apparently be transmitted from patient to patient.^4^

Whole genome sequencing (WGS) of *C. auris* isolates has indicated that there are at least four distinct geographical clades of this species; East Asia (Japan, Korea), South Asia (India, Pakistan), South Africa, and South America (Venezuela).^6^ Clades differ by tens of thousands of single nucleotide polymorphisms (SNPs) from each other, however, within each clade isolates are almost indistinguishable from each other on a DNA sequence level.^5-7^ This suggests that the *C. auris* population structure is characterized by distinct and highly variable clades that are distributed worldwide and almost non-variable clonal expansions of a single genotype within individual outbreaks.^3,4^ The origin(s) of the strong variability between and the minor variability within clades are currently unknown.

Polyploidy, aneuploidy, and gross chromosome rearrangements have been recognized drivers of genetic diversity in pathogenic and non-pathogenic fungi for some time.^8-12^ In pathogenic yeasts, such as *C. albicans,* mechanisms for ploidy shifts and chromosome rearrangements have been described, and their importance for adaptation to environmental stresses and host niches, as well as for developing resistance to antifungal drugs has been identified.^13-17^

Here, we characterize a set of 26 clinical isolates of the newly emerging human pathogenic fungus *C. auris* to understand whether its genome has undergone structural alterations potentially underlying adaptation events. This strain collection covers all four geographical clades, different levels of drug-resistance, and various sources of isolation (Table 1). All isolates were shown to be haploid, and we observed substantial karyotypical variability between strains of this fungus even when genetic diversity on a DNA sequence level had been reported to be minimal.^5-7^

**Table 1.**
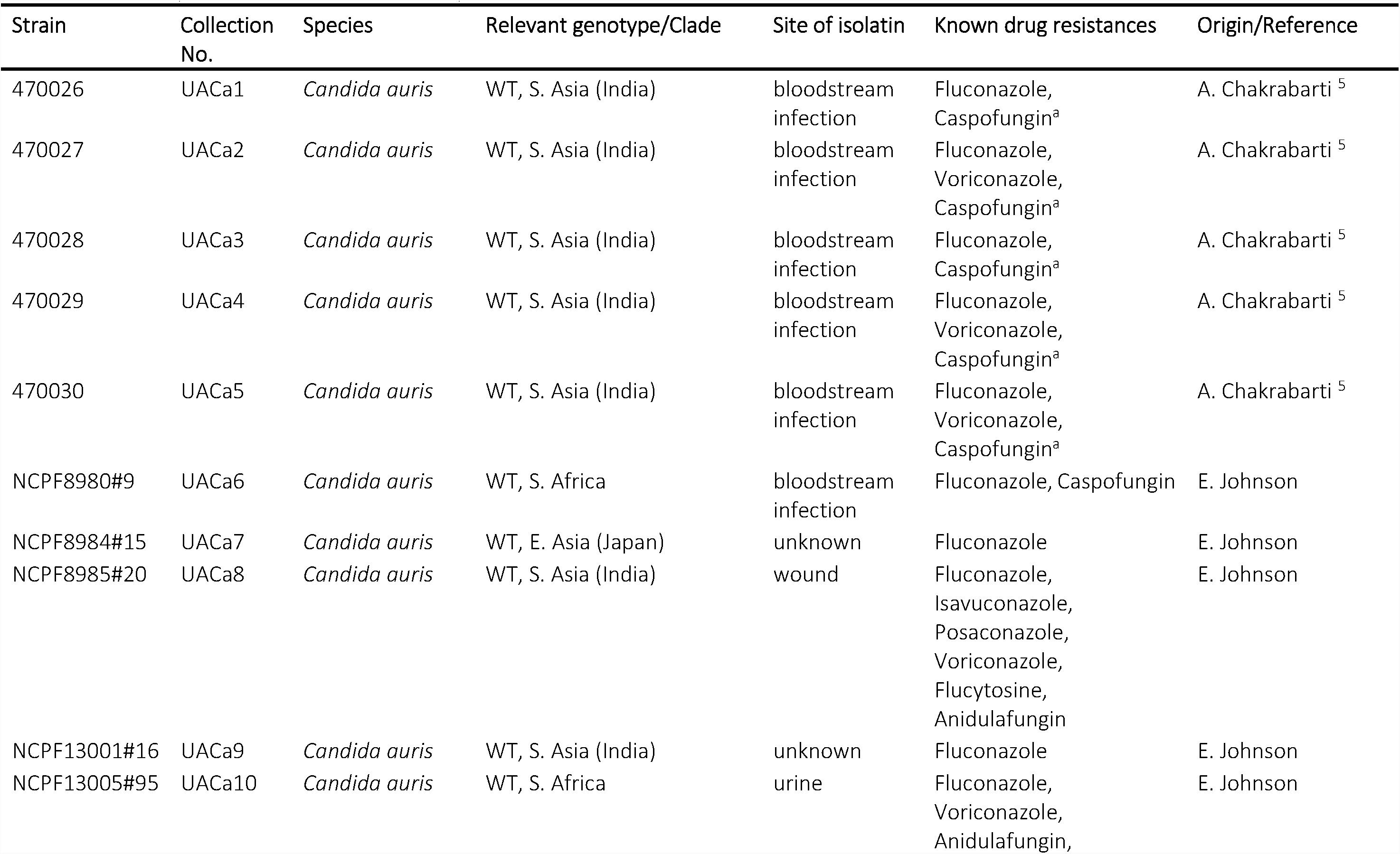
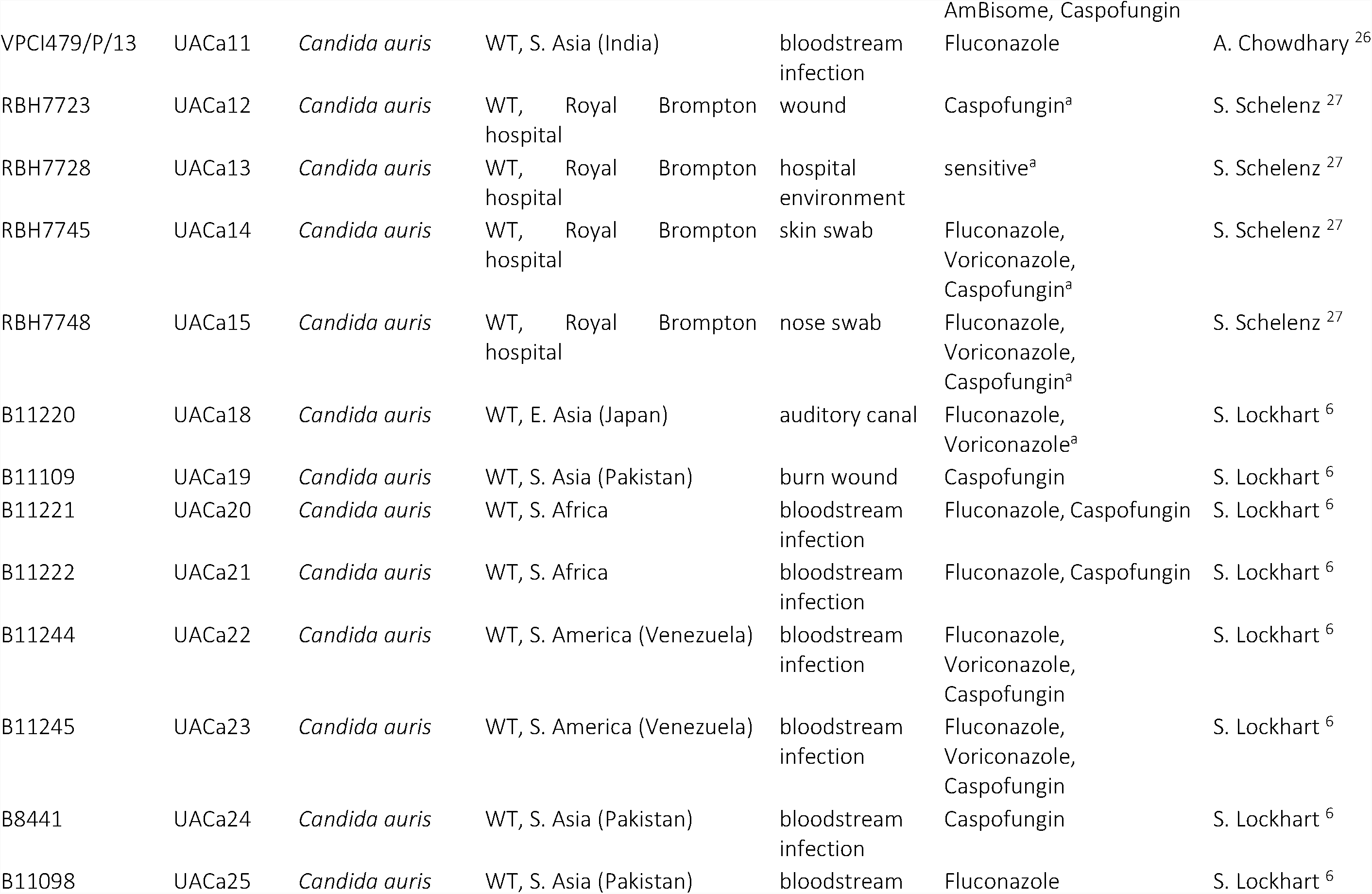
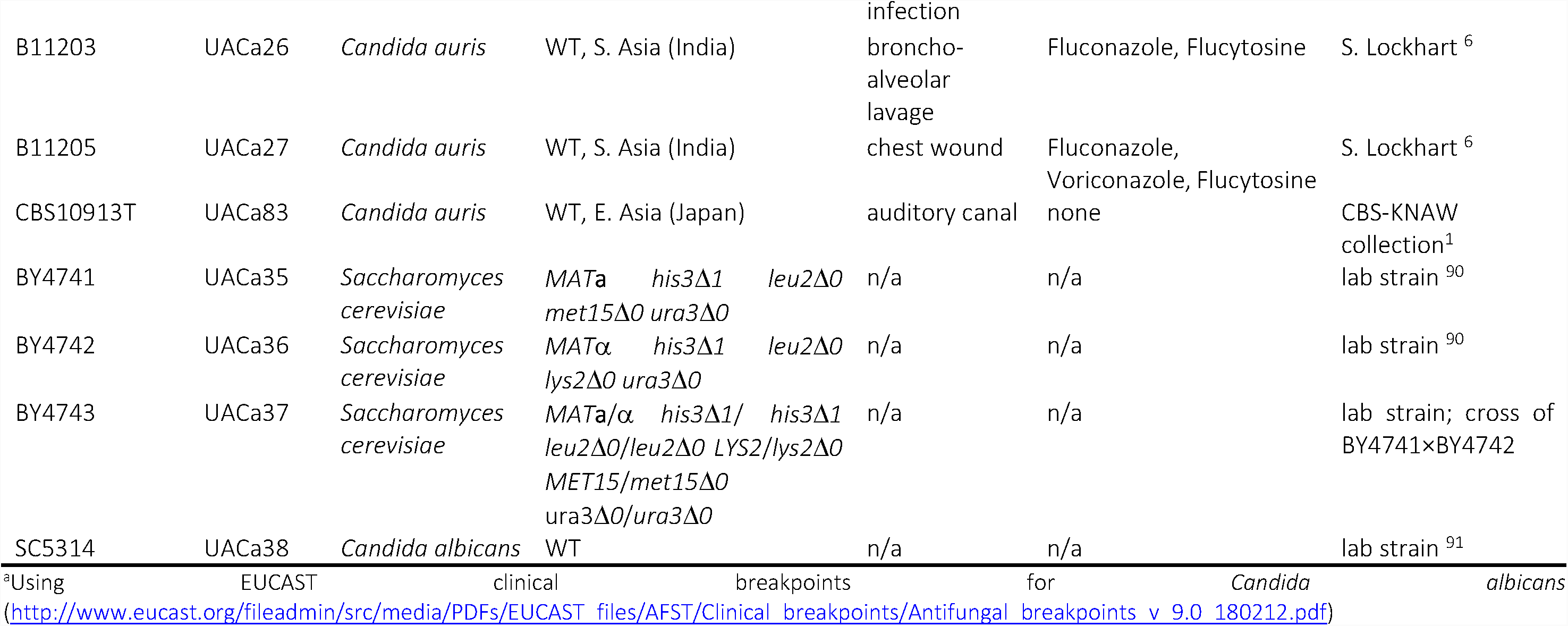
Details of yeast strains used in this study

## Materials and Methods

### Yeast strains and culture

*Candida auris* and other yeast strains used in this study are listed in Table 1. *Candida albicans* SC5314, and *Saccharomyces cerevisiae* BY4741 and BY4743 were used as control organisms. Yeast cells were grown at 30°C on YPD plates (1 % yeast extract, 2 % mycological peptone, 2 % glucose, 2 % agar; Oxoid, Basingstoke, UK) or shaking at 200 rpm in YPD broth (same as plates, but without agar).

### Flow cytometry

Processing yeast samples for flow cytometry was performed largely as previously described.^18^ Briefly, stationary-phase yeast cells were inoculated into fresh YPD broth and incubated shaking (200 rpm) at 30°C for 3 h. Cells were harvested by centrifugation (1,000 xg, 2 min), re-suspended at a concentration of 1 × 10^7^ cells/ml in ice-cold demineralized water, and fixed overnight by adding 100 % ethanol to a final concentration of 70 % ethanol. Cells were then harvested by centrifugation (1,000 x*g*, 2 min) and re-suspended in 50 mM sodium citrate (pH 7.5). After RNase A (250 μ g per 1 × 10^7^ cells) and proteinase K (1,000 μ,g per 1 × 10^7^ cells) treatment cells were transferred to 12×75 mm round bottom tubes. After adding Triton-X 100 (Sigma-Aldrich) to a final concentration of 0.25 %, and SYBR Green I (1:500; Sigma-Aldrich, St. Louis, MI, USA) as a DIMA stain, samples were incubated at 4°C overnight. Flow cytometry was performed on a BD LSR II flow cytometer (BD Biosciences, San Jose, CA, USA) using an excitation wavelength of 488 nm, SYBR Green I fluorescence was detected with a 530/30 band pass filter. Data were analyzed using FlowJo 10.2 software (FlowJo LLC, Ashland, OR, USA).

### Pulsed-field gel electrophoresis (PFGE)

Chromosomal DNA of *C. ouris* strains was embedded in agarose plugs using the CHEF Genomic DNA Plug Kit (Bio-Rad Laboratories Ltd., Hercules, CA, USA) following the instructions of the manufacturer. For some strains the cell wall digestion reaction was supplemented with Lallzyme MMX (end concentration 100 mg/ml; Lallemand Inc., Quebec, Canada). Pulsed-Field Gel Electrophoresis (PFGE) was performed on a CHEF Mapper XA System (Bio-Rad). As a standard programme *C. ouris* DNA was run for 48 h at 14°C in 1 × TAE (40 mM Tris, 20 mM acetic acid, 1 mM EDTA; pH 8.0) and 0.8 % Megabase agarose (Bio-Rad) at 3.0 V/cm applied at an 106° angle and a switch time of 500 sec. To get a better separation of smaller chromosomes *C. auris* DNA was run for 48 h at 14°C in 1 x TAE and 0.8 *%* Pulsed-Field Certified agarose (Bio-Rad) at 4.0 V/cm applied at an 120° angle, initial and final switch times of 120 sec and 240 sec using linear ramping. Gels were stained with SYBR Green I (Sigma-Aldrich) diluted 1:10,000 in 1 × TAE for at least 1 h and documented by photography under UV illumination on a Gel Doc EQ system controlled by Quantity One software (version 4.6.6) (Bio-Rad).

### Southern blot analysis

Chromosome-sized DNA bands from PFGE gels were transferred to Zeta-Probe GT membranes (Bio-Rad) by alkaline Southern blotting following previously described principles.^19^ Gels were soaked in depurfnating solution (0.25M HCl) for ∽ 25 min and, after that, in denaturing solution (1.5 M NaCl, 0.5 M NaOH) for ∽ 30 min. Using capillary transfer in denaturing solution for 24 h chromosomal DNA was immobilized on the membranes. After transfer, membranes were steeped in neutralization buffer (0.5 M Tris, pH 7.0) for 5 min, washed briefly in 2 × SSC (saline-sodium citrate; 300 mM NaCl, 30 mM NasCeHsO?, pH 7.0) and dried at room temperature.

A fragment of 836 bp was amplified by polymerase chain reaction (PCR) from the 25S rRNA region of *C. auris* strain UACa11 genomic DNA using oligonucleotides oUA367 (5’-GGCAAAACAAAGGCCGCGC-3’) and oUA368 (5’-AGTAGCTGGTTCCTGCCGAAG-3’). This fragment was used as template for a labeling PCR incorporating digoxigenin-ll-dUTP with nested primers OUA371 (5’-CCAATTCCAGGGTCACAGGCT-3’) and oUA372 (5’-CCTCAGGATAGCAGAAGCTCGT-3’) to give an rDNA probe of 759 bp (DIG DNA Labeling Mix; Roche Molecular Systems Inc., Pleasanton, CA, USA). All PCR reactions were carried out using GoTaq^®^ G2 Flexi DNA Polymerase (Promega Corp., Madison, WI, USA). Oligonucleotides were supplied by Sigma-Aldrich Co. (St. Louis, MO, USA).

Membranes were hybridized with the digoxigenin-ll-dUTP labeled rDNA probe using DIG DNA Labeling Kit (Roche Molecular Systems Inc.), and then incubated with α-digoxigenin antibody (Roche Molecular Systems Inc.) conjugated to alkaline phosphatase. Alkaline phosphatase bound to digoxigenin-ll-dUTP labeled bands was then detected on a FUSION SL Chemiluminescence Imaging System (Vilber Lourmat, Marne-la-Vall é e, France) using CPD-Star chemi-luminescent substrate (Roche Molecular Systems Inc.).

### Microevolution assay

A microevolution assay was carried out using selected strains from each clade (UACa11, UACa18, UACa20, and UACa22) to test whether karyotype variation can be induced by particular growth conditions: strains where separately passaged five times through YPD broth at 30°C (control), YPD broth at 42°C (heat stress), synthetic defined liquid medium containing 2 *%* sorbose (SSD) (6.7 g/l yeast nitrogen base with amino acids, Sigma-Aldrich) at 30°C (osmotic stress), and YPD broth containing 100 mM hydroxyurea (HU; Formedium, Norfolk UK) at 30°C (DNA replication stress). All passages were performed using the same experimental scheme. A passage consisted of (1) growing cells in liquid culture overnight, (2) plating 100-200 cells on appropriate solid medium until single colonies were visible (2-3 days, except on 2 % sorbose where incubation took 1 week), (3) selecting the five largest single colonies, (4) suspending each colony in 1 ml of sterile water and making four 1:10 serial dilutions to test resulting isolates in spot assays, (5) selecting the three fastest-growing spot to further analysis and the fastest one to inoculate a fresh overnight liquid culture starting the next passage.

Isolates from the first and the fifth passage of each strain and condition were subjected to PFGE analysis (see above), and spot assays under the same conditions used for each passage (except for DNA replication stress where isolates were also tested on plates containing 200 mM HU); the parental strain was always included for comparison. For those spot assays, isolates and parental strains were grown in liquid culture overnight, cell concentrations were determined by measuring optical density of the culture at a wavelength of 600nm (OD_600_) on an Ultraspec 2000 (Pharmacia Biotech, Sweden) spectrometer. Previous calibration defined a *C. auris* culture of OD600 = 1 to contain 3×l0^7^ cells/ml. Dilutions were made for spots to consist of 10^2^, 10^3^, 10^4^, and 10^5^ cells. Appropriate plates were grown for 1-6 days depending on conditions and temperature used for each passage.

## Results

### *Candida auris* is a haploid fungus

Polyploidy and complex aneuploidy play a major role in the capability of fungal pathogens to adapt to various stresses and to the changing condition within host niches.^10,11^ These overall genome shifts in chromosome number have been characterized as drivers of increased genetic diversity in *Cryptococcus neoformans, Candida albicans,* and also for *Candida lusitaniae*-a close relative of *C. auris*.^20-25^

We were, therefore, interested in determining the ploidy of clinical *C. auris* isolates to understand whether chromosome number variations and whole genome duplication potentially are adaptive strategies employed by *C. auris.* In total 25 *C*. *auris* strains covering all four geographical clades, comprising antifungal-sensitive,-resistant, and-multiresistant isolates from various sources of infection, as well as strains from a single outbreak (Royal Brompton hospital, London, UK) (Table 1) were tested by flow cytometry.^5,6,26,27^ Haploid and diploid *Saccharomyces cerevisiae* strains and the diploid *C. albicans* laboratory strain SC5314 were used as references. Estimates from whole genome sequencing suggested that *C. auris* has a similar genome size as *S. cerevisiae* at approx. 12 Mbp.^6,28^ As expected, C. *albicans* SC5314 had a similar cell cycle profile as a diploid *S. cerevisiae* strain (Fig. 1). In contrast, the genome size of all 25 *C. auris* strains was found to be consistent with containing a haploid chromosome complement (Fig. 1), although the resolution of flow cytometry probably does not allow us to unequivocally exclude the occasional disomy of one of the smaller chromosomes. However, in none of the 25 strains disomies were apparent in the karyotype analysis (see below).

**Figure 1.**
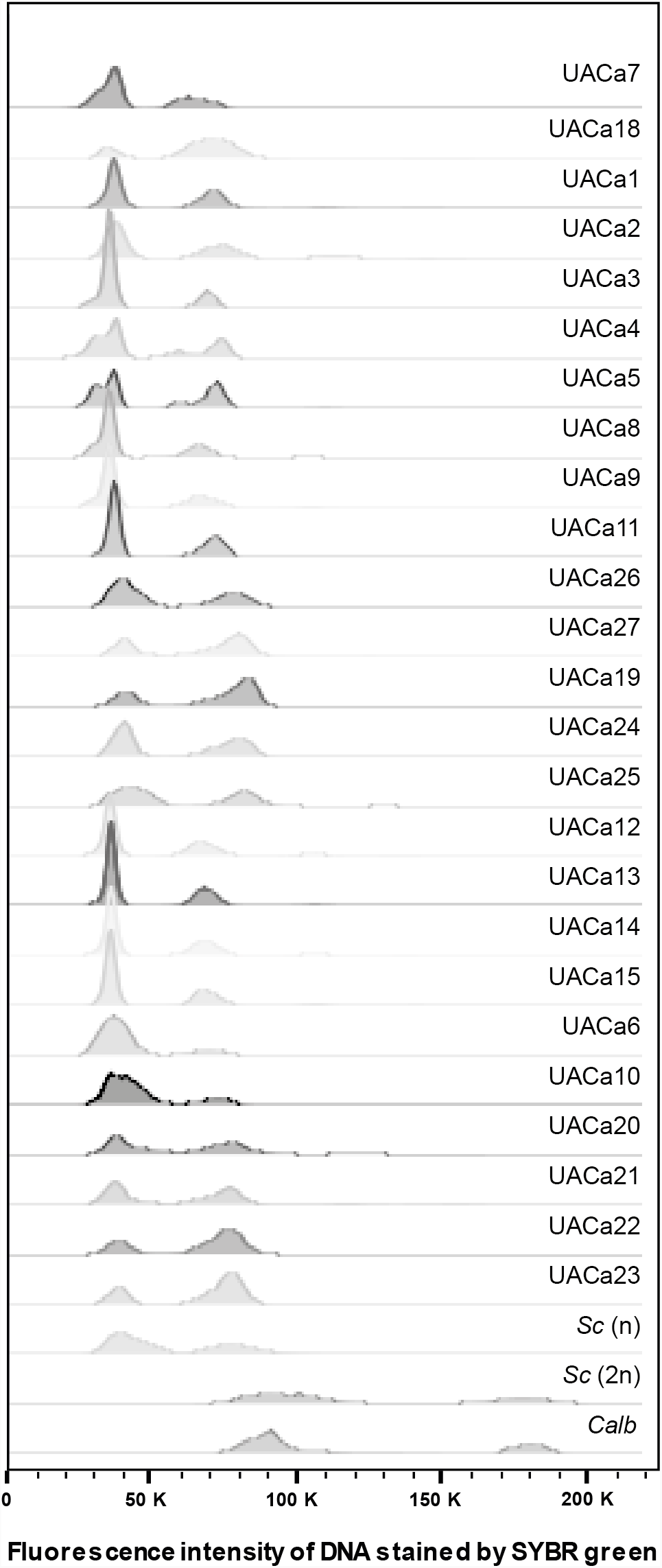
Cell cycle profile of clinical *Candida auris* isolates. Histogram showing cell cycle profile obtained by flow cytometry after staining DNA with SYBR green of 25 *C. auris* strains (Table 1). Diploid *Candida albicans (Calb)* strain SC5314, haploid (BY4741) and diploid (BY4743) strains of *Saccharomyces cerevisiae* (5c) were incuded as controls. C. *auris* has the same cell cycle profile as haploid *S. cerevisiae, C. auris* strains are grouped according to their taxonomical position within the 4 geographical clades: E. Asia (UACa7 & UACa18), S. Asia-lndia (UACal-5, UACa8, UACa9, UACa11, UACa26-27), S. Asia-Pakistan (UACal9, UACa24-25), strains from the Royal Brompton hospital outbreak (UACal2-15), S. Africa (UACa6, UACalO, UACa20-21), and S. America (UACa22-23).

This result indicates, that major ploidy changes do not appear to be a mechanism *C. auris* employs to adapt to environmental challenges. If it does, this must be only temporarily with diploids, polyploids, or aneuploids returning quickly to a haploid stage after these challenges are removed.

### *Candida auris* clinical isolates have a plastic karyotype

Since all the *C. auris* isolates tested turned out to be haploid (or near-haploid) we wondered whether its genetic diversity might be generated by gross chromosome rearrangements. To test this we utilized <u>P</u>ulsed-<u>F</u>ield <u>G</u>el <u>E</u>lectrophoresis (PFGE) to separate *C. auris* chromosomes (Fig.2). We characterized the same 25 strains covering all four geographical clades and the single hospital outbreak, and an additional East Asian isolate, UACa83 (the type-strain of C. auris, CBS10913T) (Table I),^5,6,26,27^ *C. auris* isolates show chromosome numbers from five to seven, ranging from ∼0.7 Mbp to ∼3.25 Mbp in size (Figs. 2,3); chromosomes will be referred to by their size. Additionally, we probed the karyotypes for the location of the repetitive rRNA gene clusters (rDNA) by Southern blotting (Figs. 2, 3). It should be noted that chromosomal bands of similar size are difficult to separate using the PFGE system. However, we have considered the presence of two chromosomes when the band intensity is clearly higher, e.g. chromosomal band around 1 Mbp in UACa20 likely or chromosomal band around 1.35 Mbp in UACalS possibly each contain 2 chromosomes.

**Figure 2.**
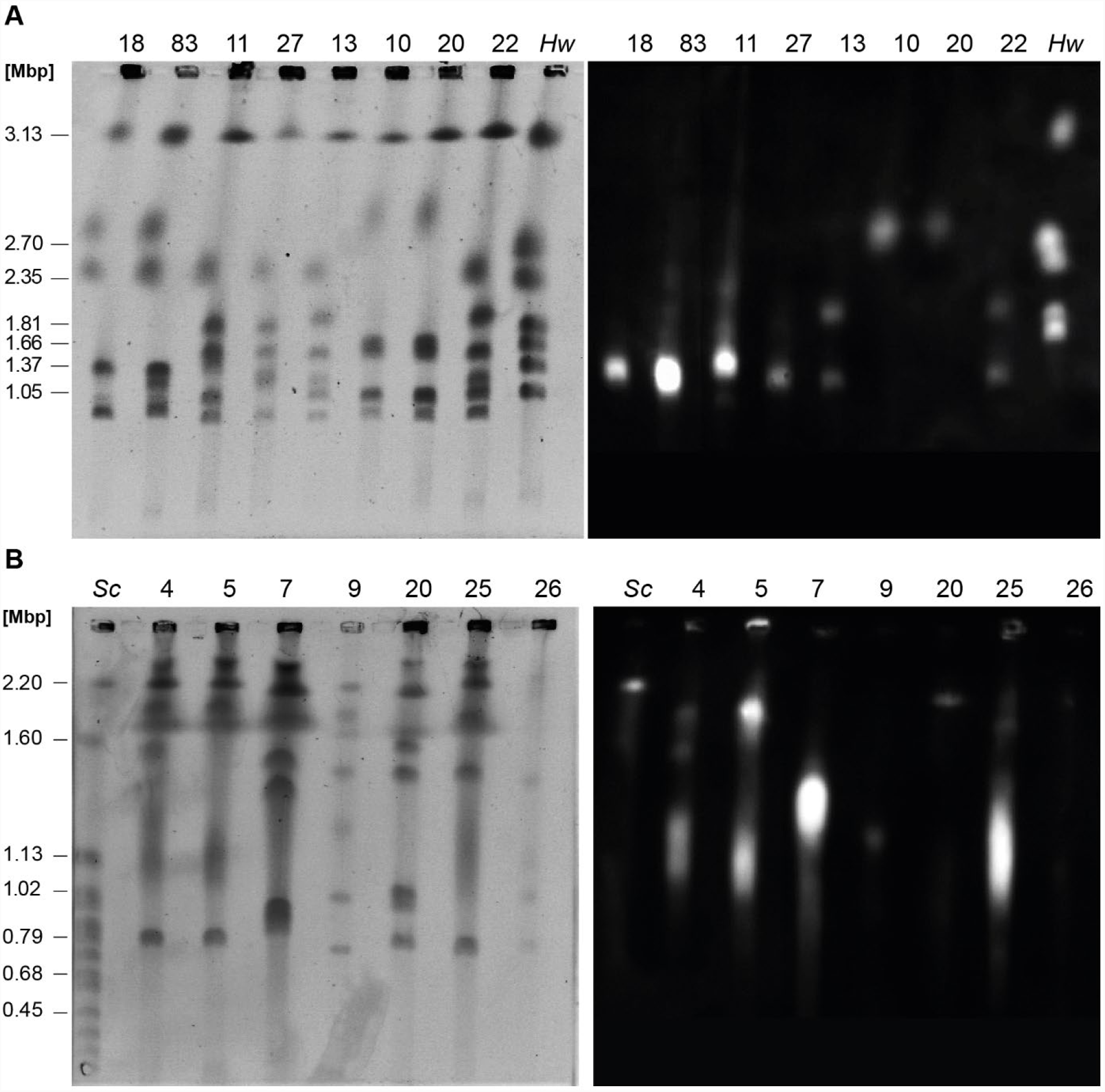
Representative PFGE karyotypes of clinical *Candida auris* isolates. PFGE karyotypes (left panels) and associated Southern blots using a rDNA probe to detect rRNA gene clusters (right panels) of the indicated strains representing examples of E. Asian (UACa7, UACa18, UACa83), S. Asian (UACa4, UACa5, UACa9, UACa11, UACa25, UACa26, UACa27), S. African (UACalO, UACa20), and S. American (UACa22) clades, as well as one isolate from the outbreak at the Royal Brompton Hospital, London, UK (UACal3) (Table 1). **(A)** Gel run at standard conditions (0.8 *%* Megabase agarose, 1 × TAE, 48 h, 14°C, 3.0 V/cm, 106°, switch time 500 sec), *Hansenula wingei (Hw)* CHEF DNA size marker (Bio-Rad) serving as standard (size of chromosomal bands in Mbp indicated on the left). **(B)** Gel run at conditions to resolve smaller chromosomes (0.8 % Pulsed Field Certified agarose, 1 × TAE, 48 h, 14°C, 4.0 V/cm, 120°, switch times: linear ramping 120-240 sec), *Saccharomyces cerevisiae* (Sc) CHEF DNA size marker (Bio-Rad) serving as standard (size of some chromosomal bands in Mbp indicated on the left).

**Figure 3.**
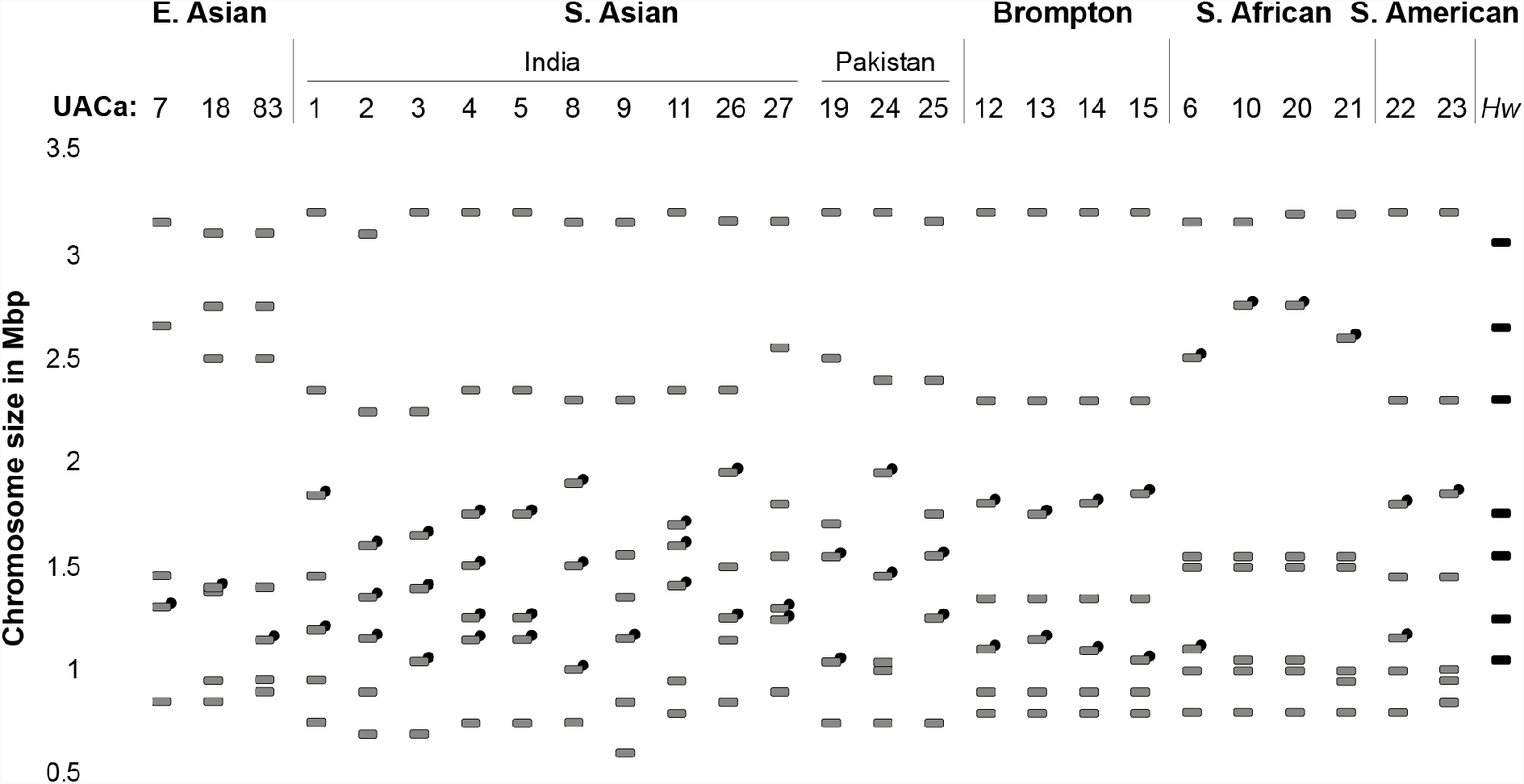
Schematic of karyotypes of 26 clinical *Candida auris* isolates. Strains are grouped according to their taxonomical position within the four geographical clades: E. Asia, S. Asia (India and Pakistan subgroups indicated), S. Africa, and S. America. The four strains isolated from the outbreak in the Royal Brompton Hospital, London, UK (Brompton) are indicated as a separated group. Chromosome sizes were measured using the *Hansenula wingei (Hw)* CHEF DNA size marker (Bio-Rad) and strain UACa11 as standards, both of which were run alongside the samples in each gel. Chromosomes are represented as light-grey rectangles for different strains or black rectangle for the *Hw* marker. Chromosomes harboring rRNA gene clusters are indicated by a black circle (·).

The four strains from the Royal Brompton hospital outbreak (UACal2-UACal5) had similar karyotypes to each other with seven detectable chromosomal bands. The two chromosomes bearing rDNA repeats displayed subtle size differences between these particular set of strains (Fig. 3). This is not unusual in fungal pathogens, and similar observations have been made in other *Candida* species.^29-32^ Previously reported electrophoretic karyotypes of clinical C. *auris* isolates from three different hospitals in Korea appeared fairly uniform displaying five or six chromosomal bands, but revealed subtle differences after restriction endonuclease digest with the rare cutting enzyme *Notl*.^33^ This is similar to the E. Asian strains of this study, which apparently have five to seven chromosomes one of which carries rRNA gene repeats. The S. African and S. American isolates we studied all had seven chromosomes and with the exceptions of UACa6 (S. Africa) and UACa22 (S. America) only one of the chromosomes was rDNA-bearing (Fig. 3). In contrast, chromosome numbers in S. Asian isolates ranged from six to seven, and, except for UACa9, all S. Asian strains had at least two chromosomes carrying rRNA gene repeats (Fig. 3). The range of chromosome numbers and of chromosome size distributions in S. Asian isolates maybe reflects the comparatively large intra-clade genetic diversity of this strain cluster^6^, but we cannot exclude this to be an issue of having only a small sample sets available for non-S. Asian isolates. Interestingly, the four strains from the Royal Brompton hospital outbreak with their seven chromosomes, two of which are rDNA-bearing, appeared similar to S. Asian strains, which they are also most closely related to according to whole-genome sequence analysis.^7^

Adding estimated chromosome size estimates from PFGE points towards a range of genome sizes between ∼10 Mbp and ∼13 Mbp which is a reasonable fit to the 12.5 Mbp suggested by whole-genome sequencing.^6^ Complete assembly of whole-genome sequences into chromosome-sized contigs to create physical maps of *C. auris* genomes will allow full appreciation of the genome structure of this fungus. Indeed, a recent study reported seven contigs for two C. *auris* isolates, in our PFGE analysis the corresponding strains UACa20 (B11221) and UACa24 (B8441) also display seven chromosomal bands.^34^

Electrophoretic karyotyping revealed that *C. auris* isolates differed considerably in chromosome numbers and sizes, both within and between geographical clades. This plasticity was somewhat unexpected considering the genetic uniformity of *C. auris* on a DNA sequence level within geographical clades and within hospital outbreaks, and indicates that gross chromosome rearrangements might be a mechanism *C. auris* employs to generate genetic diversity during adaptation to environmental challenges.^6,7,35^

### *C. auris* undergoes karyotype rearrangements in stress conditions

In order to get insight about *C. auris* fitness and its relation to the karyotype variation observed in different clinical isolated, four strains (UACa11, UACa18, UACa20, and UACa22), one from each clade (Table 1), were selected to undergo a microevolution experiment. Strains were grown under four different conditions through five passages each (see Materials and Methods): standard YPD broth (control); heat stress at 42°C; osmotic stress using 2 % sorbose, which mimics the effect that echinocandin-type antifungals have on cells; and DNA replication stress using hydroxyurea (HU), an inhibitor of the enzyme ribonucleotide reductase (RNR) which depletes nucleotide pools.^36,37^ These conditions have been described previously as factors inducing genome instability in fungi. In *C. albicans,* treatment with 2 *%* sorbose is a classic example for inducing changes in the karyotype, which offer a phenotypic advantage in this stress.^38,39^ Several karyotype changes have been observed in this condition affecting different chromosomes. Specifically, extra copies of chromosome III containing *SOU1,* a key gene for the ability to use L-sorbose, or loss of one copy of chromosome V, containing *CSU1* an inhibitor of *SOU1* have been observed.^40^ Introduction of heat stress has been shown to induce ploidy variation in *C. albicans* and other fungal species, this is associated to the functions of the Hsp90 chaperone complex.^41-42^ Replication stress, for example by treatment with HU, has been associated with genome instability in a wide range of organisms, including yeast and human cells.^43-44^

In our microevolution study karyotype modifications including appearance, disappearance, or size changes of different chromosomal bands were observed in all tested strains. However, the frequency and type of modification was different depending on the strain and condition used (Fig. 4).

**Figure 4.**
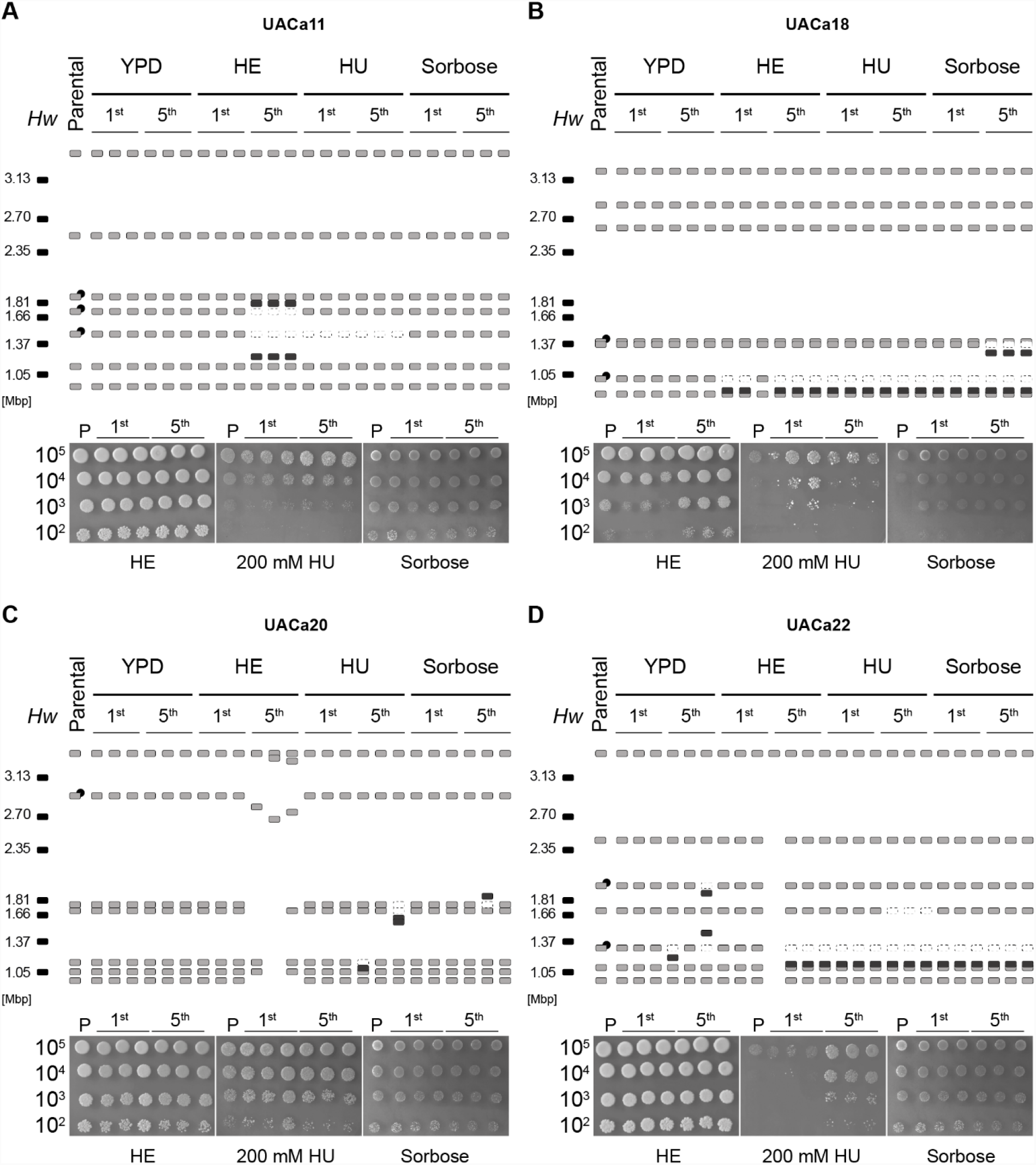
Karyotype variation during microevolution of *Candida auris* isolates. Schematic representation of karyotypes (top panels) and spot assays (bottom panels) of four *C. auris* isolates covering all four established clades **(A)** UACa11 (S. Asia), **(B)** UACa18 (E. Asia), **(C)** UACa20 (S. Africa), and **(D)** UACa22 (S. America)-obtained after the 1^st^ and 5^th^ passages in four different growth conditions: YPD at 30°C; YPD at 42°C, heat stress (HE); YPD containing 100 mM hydroxyurea at 30°C (HU); and *2%* sorbose in synthetic defined medium at 30°C (sorbose). Chromosomes are represented as light-grey rectangles for different strains or black rectangle for *Hansenula wingei* (*Hw)* CHEF DNA size marker (Bio-Rad) used as a chromosome size standard (numbers represent size in Mbp). Chromosomal bands that seemingly disappear in comparison to parental strains are represented as empty rectangles with dotted lines. Dark-grey rectangles represent new chromosomal bands appearing in comparison to parental strains. In strain UACa20 under heat stress the karyotype changes observed indicate massive chromosome rearrangements. Chromosomes harboring rRNA gene clusters are indicated by a black circle (·) only in parental strains. Spot assays show the parental strain (P) and derived isolates for comparison, grown in the same conditions used for the respective passages, except for HU for which a higher concentration (200 mM) than during the passages (100 mM) is used. Serial dilutions contain 10^5^, 10^4^,10^3^, and 10^2^ cells. Isolates were grown for 1 day in heat stress *(42°C),* 3 days in HU (due to slow growth, UACa18 and its derivatives were grown for 6 days), or 3 days in 2 % sorbose.

The S. Asian strain UACa11 has shown chromosome modification under heat and DNA replication (HU) stress conditions, always related to chromosomes carrying rDNA repeats (Fig. 4A). In HU, the 1.35 Mbp chromosomal band seemingly disappears after only one passage. In heat stress, the 1.35 Mbp and 1.6 Mbp chromosomal bands disappeared, and two additional ones appeared (around 1.1 Mbp and 1.7 Mbp), most likely due to loss and gain of DNA from the corresponding original chromosomes.

The E. Asian isolate UACa18 showed a tendency to lose the 0.95 Mbp chromosomal band in all stress conditions, but not when growing in YPD at 30°C (control), most likely changing the size to a slightly smaller chromosome of around 0.9 Mbp (Fig. 4B). When treated with sorbose, a new chromosome of around 1.3 Mbp appeared after five passages in UACa18, likely because of the reduction in size of one of the two 1.35 Mbp chromosomes.

Looking at the S. African strain UACa20, only minor changes were observed after five passages in sorbose and HU (Fig. 4C). Specifically, one alteration was observed in sorbose where one of the isolates seemed to gain DNA in the 1.65 Mbp chromosome, andinHUwherethel.05 Mbp chromosome decreased in size in one isolate, and two other chromosomes (1.6 and 1.65 Mbp) seemingly reduced size in another isolate. However, under heat stress conditions drastic changes were observed, the number of chromosomes reduced from seven to six, five, or three bands in three different isolates.

Finally, the S. American strain UACa22 was the most plastic, showing karyotype changes even when grown in control conditions (YPD, 30°C), in this case chromosomes carrying the rDNA tend to change size, losing or gaining DNA (Fig. 4D). In almost all conditions the 1.2 Mbp chromosome tends to reduce in size. Besides, a second chromosome (1.6 Mb) was seemingly lost in the presence of HU after five passages.

Despite the karyotype alterations observed, a moderate improvement in growth was detected only in a few cases (Fig. 4): isolates from UACa18 microevolved in 2 % sorbose from the first passage, and two isolates on 200 mM HU after one passage in 100 mM HU; and isolates from UACa22 on 200 mM HU for five passages of 100 mM HU. Although these improvements in growth could be correlated with changes in the karyotype in some cases (see above). This clearly is not probable for all the cases, e.g. isolates of strain UACa20 which grow better in sorbose, but did not show any difference in karyotype in comparison to the parental strain, and modifications in karyotype which did not obviously offer any improved fitness in the condition tested. Interestingly, the massive karyotype variation observed in microevolved UACa20 isolated obtained under heat stress did not show an apparent difference growing at 42°C compared to the parental strain. Isolates from the strain UACa11 did not show improvement in growth under any conditions.

In our microevolution study we tested whether under heat, osmotic, or DNA replication stress karyotype changes are induced in *C. auris,* similar to other fungi.^45-46^ The frequency of changes is apparently higher in stress conditions, but minor alterations could also appear when *C. auris* was grown in rich medium at 30°C. We also observed that these changes can be associated with fitness benefits in some cases. However, not all changes provide an advantage, indicating that the karyotype modifications we observed could be stochastic, as previously reported for other fungi.^47-48^ Strikingly, the strain UACa20 showed massive chromosome rearrangements reducing a seven chromosome karyotype down to three chromosomes in one isolate, demonstrating that drastic modifications of the genome structure do not necessarily impinge on viability in *C. auris;* this might provide opportunities for general fitness adaptation.

## Discussion

Here we show that the genome of *C. auris* undergoes substantial karyoytypic reorganiztion under stress conditions (Fig. 4), similar to other related fungi.^45,49,50^ Karyotype variation between different strains of the same species is common in fungi, and it has been proposed as a quick solution for adaptation to environmental changes. This variation of genome organization can take on different expressions, e.g. ploidy variation, chromosome copy number variation, or gain and loss of supernumerary chromosomes.^10,46,51-54^ High levels of genetic diversity can be introduced into a population by changes in ploidy. In yeast most of these changes produce a mixture of aneuploid populations offering a rapid solution for stress adaptation, this has been described as the norm for fungi.^10^ These changes in genome structure might be introduced by various mechanisms usually related to mitosis, sexual or parasexual reproduction. Although, once aneuploidy arises in *S. cerevisiae,* rates of chromosome loss, genetic mutation, and microsatellite instability increases, usually leading to proliferative disadvantages.^20,55-58^ Having said that, aneuploid isolates of *S. cerevisiae* exposed to a range of stress conditions display a growth advantage, when the stress condition is maintained.^59-61^ For that reason, aneuploidy is often lost, when the stress is eliminated.^40^ Aneuploidy changes are more likely in diploid strains where a second copy of the genome can be used to maintain viability, and to potentially restore the original euploid state later on. Indeed, diploid cells can tolerate a more diverse repertory of rearrangements than haploids, and haploid *S. cerevisiae* strains are genetically more stable showing lower rates of chromosomal rearrangements than diploid cells.^62-64^ Since *C. auris* is haploid (Fig. 1), and so far, neither polyploid states nor sexual reproduction have been described, we hypothesize that several of the mechanisms generating genome diversity in many fungi might be defective in *C. auris* at least to some degree. Further studies to elucidate the life cycle of C. *auris* will be necessary to shed light on this issue.

As a haploid species, the variation observed in *C. auris* would most likely be due to gross chromosome rearrangements, and/or possibly copy number variation (CNV) events of chromosomal sections. CNVs are a frequent reason for changes in the genome organization of haploid fungi.^50,64^ CNVs of a yet to be identified genomic regions, potentially repetitive, could explain the changes in size observed in the karyotype of *C. auris,* e.g. strain UACa20 after heat stress, the 0.95 Mbp chromosomal band in UACa18, or 1.2 Mbp chromosome in UACa22 (Fig. 4). Intriguingly, the appearance of CNVs increases environmental adaptation, for example increased resistance to antifungal azoles in *C. olbicons.*^*^65-68^*^ In *C. glabroto* most frequent changes in karyotype are caused by chromosomal translocations and CNVs (including CNVs at tandem gene repeats).^69^ *C. glabrata* might be more prone to CNVs and chromosome rearrangements than *S. cerevisiae,* because it contains more minisatellites and megasatellites (giant minisatellites with lengths up to 10 kb).^70,71^ In general, repetitive regions are known as a principal reason for CNVs and gross chromosome rearrangements in yeasts, such as transposons (Ty elements and solo LTR elements in *S. cerevisiae),* telomeres, or rDNA.^64,72-77^ One of the best-studied repetitive elements is the rDNA. The rDNA locus is highly conserved in eukaryotes, with a large number of tandemly repeated sequences, comprising both genes and intergenic regions with noncoding elements. In 5. *cerevisiae* the rDNA consists of ∼ 150 tandem copies of a 9.1-kb sequence encoding the ribosomal RNAs similarly to *C. albicans,* in which the haploid genome encodes a single ∼ 12 kbp rDNA region located in the chromosome R.^78,79^ Repetitive in nature, the rDNA is highly recombinogenic, enabling fluctuations in rDNA copy numbers, related to the loss of global chromosomal stability in *S. cerevisiae,* especially during senescence.^80,81^ Furthermore, in C. *albicans* chromosome R has been described as more unstable than the other chromosomes within the complement, and thus more frequently displaying size changes.^82,83^ Indeed, our set of strains from the outbreak at the Royal Brompton hospital shows moderate size changes in the chromosomes harboring the rDNA repeats only (Fig. 3). In the first draft genome obtained for *C. auris* strain Ci6684, seven rRNA gene repeats were described, since this is an incomplete assembly containing 99 scaffolds, the true number of rDNA loci will likely be smaller.^84^ Our survey of PFGE karyotype by Southern blotting using *C. auris* rDNA as a probe indicates that between one and four chromosomes harbor an rRNA gene repeat (Fig. 3), which clearly could be the source for some of the rearrangements we observed (Figs. 3 & 4). The presence and involvement in chromosome rearrangements and CNVs of other repetitive elements in *C. auris,* such as retrotransposon or minisatellites will thus be an interesting aim for future study. In our microevolution study, we also observed the disappearance of chromosomal bands, which cannot obviously be explained by a change in size, like in strain UACa11 and UACa22 in HU (Fig. 4). In these instances it is likely that a change in size occurred, and the new size is being masked by another chromosomal band. Due to *C. auris* being haploid, we can exclude a complete loss of a chromosome, as has been described in diploid *C. albicans.*^*^45,66^*^ However, other possibilities should be taken into consideration, e.g. the presence of supernumerary chromosomes. Especially, in fungal plant pathogens supernumerary chromosomes often confer an adaptive advantage in particular habitats or on certain hosts.^53^ For example the opportunistic pathogen *Fusarium oxysporum,* which can infect plants and animals, supernumerary chromosomes have been described as key determinants in its broad host range, while the conserved core genome maintains essential house-keeping functions.^54^ The appearance of a novel chromosomal band (1.3 Mb) has been observed in the strain UACa18 after passaging through 2 % sorbose (Fig. 4). There are several potential explanations for this observation, among the more likely ones are, (I) that the chromosomal band at 1.37 Mbp contains two chromosomes, (II) that in the microevolved UACa18-derivative the resulting population represents a mixture containing cells with the original rDNA-bearing 1.37 Mbp chromosome and cells with a considerably shorter version of this chromosome, or (III) similar to *C. glabrata,* that the appearance of the novel chromosome originated from segmental duplications in one of the two smaller chromosomes (Fig. 4).^71,85^ These novel chromosomes in *C. glabrata* carry duplicated genes potentially involved in yeast-host interaction and virulence, including the ABC transporter family genes, which play a role in multidrug resistance.^71,85^

Strikingly, we observed massive chromosome rearrangements in the strain UACa20 during our microevolution assay under heat stress reducing the number of chromosomes from seven down to three chromosomes in one isolate (Fig. 4). In fungi, two mechanisms have been suggested as a cause for reduction in chromosome numbers, telomere-to-telomere fusions and inactivation of one centromere, or breakage of a chromosome at a centromere and posterior fusion to telomeres of other chromosomes, being the first one the most frequent.^86,87^ Viable strains of S. *cerevisiae* with a genome consisting of only one or two chromosome can be obtained by CRISPR-Cas9-mediated engineering of end-to-end chromosome fusions and centromere deletions, though these strains display a somewhat reduced fitness.^88,89^ The strains obtained for *C. auris* in this study with three, four or five chromosomes, instead of seven in the parental, did not show any obvious growth defects, and likely are thus fully viable (Fig. 4). Although changes in the karyotype of *C. auris* are not obviously faster than in any other related fungus, our results demonstrate that it is capable of undergoing and maintaining drastic alterations of its genome structure. This could be a source of adaptation to stressful conditions, and could underpin the virulence of this dangerous fungus.

## Acknowledgements

We are grateful to Arunaloke Chakrabarti, Anuradha Chowdhary, Elizabeth Johnson (PHE), Takashi Kubota, and Shawn Lockhart (CDC) for providing strains. Flow cytometry was performed at the Iain Fraser Cytometry Centre (IFCC), University of Aberdeen (Raif Yuecel). This work was funded by the Medical Research Council (MRC) Centre for Medical Mycology at the University of Aberdeen (MR/P501955/1), a Wellcome Trust Institutional Strategic Support Fund grant awarded to the University of Aberdeen (204815/Z/16/Z), a Tenovus Scotland project grant (G17.02), a Royal Society Research Grant (RG140254) to AL, and Wellcome Trust Strategic Award, Senior Investigator and Collaborative Awards (080088, 086827, 075470, 099215, and 097377) to NARG,

